# Angiogenesis inhibiting capacity of *Basella rubra* and *Syszygium cumini* fruit extracts using chorioallantoic membrane assay

**DOI:** 10.1101/201566

**Authors:** Hannah Paula G. Galang, Milton Norman D. Medina, Pedrito M. Castillo

## Abstract

Angiogenesis is a physiological process of new blood vessel development from pre-existing capillaries. This process is also the main qualification for tumor growth and plays a vital role in tumor invasion and metastasis. Nevertheless, angiogenesis inhibitors can be used to impede abnormal blood vessel growth. The study was conducted to assess angiogenesis inhibiting potential of *B. rubra* (Alugbati) and *S. cumini* (Lumboy) fruit extracts as cell mass growth retardants using Chorioallantoic Membrane (CAM) Assay in *Anas platyrhyncho* (common Duck) embryo. The treatments were individually compared to Retinol palmitate as positive control and 0.9% sterilized normal saline solution as negative control. Data were gathered and analyzed using Analysis of Variance (ANOVA) and Least Significant Difference (LSD) to test for the significant anti-angiogenic activity and pair-wise comparison among the treatments respectively using a p-value of less than 0.01. Analysis showed that there is no significant difference between the fruit extracts and positive control while a significant difference between the fruit extracts and negative control was observed. Both treatments showed very good anti-angiogenic effect with average scores of 1.33 and 1.67 respectively compared to the positive control with a scoring averaging of 1.78. Furthermore, toxicity test is recommended for both treatments.

## INTRODUCTION

Angiogenesis or the growth of new blood vessels from pre-existing blood vessels is a hallmark of tumor development, metastasis, and other pathologic conditions (Hillen & Griffioen, 2007).If the balance is disturbed, there will either be too much or too little angiogenesis that could lead to several diseases such as skin diseases, stroke, and pregnancy related diseases like intrautine growth restriction and preeclampsia (Chung, Lee & Ferrara, 2010). The process of angiogenesis and its diverse signaling pathways remarkably provide various options for therapeutic intervention (Ahmed & Bicknell, 2009).Recent findings reveal that cancerous tumor cannot grow to more than 2mm in diameter unless there is development of new blood vessels thus, showing the essential role of angiogenesis in tumor growth and invasion(Nishida, et al., 2006).This leads to the development of anti-angiogenic drugs, a major area of oncologic research today(Carmeliet, 2000).

Vasculature in angiogenesis can be started abnormally to generate new blood vessels during pathological conditions such as cancer which is the third leading cause of morbidity and mortality in the Philippines (Ngelangel & Wang, 2015). There are 14.1 million cancer cases around the world in 2012 and is expected to increase to 24 million by 2035 (Ferlay, et al. 2015).

Recent research found that fruits of twining herbaceous vine *B. rubra* (Alugbati)and *S.cumini* (Lumboy)were anthocyanin-rich fruit (Kumar, et al., 2013; Banerjee, 2015). Several researches revealed that anthocyanins are potential anti-cancer compounds (Joshi & Goyal, 2011; Sugata, Lin, & Shih, 2015) which is abundant in almost all types of vascular plants (Medina et al. 2016). Hence, this study utilized the anthocyanin content of both fruits as retardants of abnormal cell mass growth restraining formation of new blood vessels using Chorioallantoic Membrane (CAM) Assay on duck’s embryo.

The fruit extracts are expected to lessen the abnormal cell mass growth which signifies a priority area in vascular tumor biology and could give insights to the importance of anti-angiogenesis in treating and constraining the development of several other diseases (Carmeliet, 2008). The present paper showed the results of the experiment on angiogenesis inhibiting capacity of *B. rubra* (Alugbati) and *S. cumini* (Lumboy) fruit extracts using chorioallantoic membrane assay.

## METHODOLOGY

### Research Design

There were four (4) treatments used with three (3) trials with three (3) replicates were prepared for each treatments. The following treatments were: *B. rubra* (Alugbati), *S. cumini* (Lumboy), Retinol palmitate (Afaxin, 25,000IU) as positive control and 0.9% sterilized normal saline solution as negative control. Nine (9) duck eggs were used in each treatment.

### Preparation of Plant Material

Fresh *B. rubra* (Alugbati) and *S. cumini* (Lumboy) fruits were properly cleaned using running tap water and then air-dried. The fruits were placed in a sterile container and soaked in sterile distilled water for 24 hours. After soaking, the fruits were blended and then filtered using sterile filter cloth. The extracts were placed in the refrigerator at 2°C until used.

### Chorioallantoic Membrane (CAM) Assay

The method of Marchesan et al. (1998) was used with modification. Freshly laid duck eggs were bought from a commercial handler and immediately incubated at *37*°C for 7 days with constant agitation. The eggs were properly arranged, randomized, and labeled in the tray. After incubation, the eggs were opened at the flattened base for removal of 5 ml albumen and sealed with parafilm. The eggs were then horizontally arranged and 1-2 cm in diameter hole was opened on the snub side to expose the chorioallantoic membrane of the duck egg. The eggs were then sealed using laboratory films and incubated until the addition of treatment.

The prepared treatments were added to the duck eggs. Each egg was dispensed with 200μl of the treatment on the air sac. The eggs were then incubated for another 5 days. This was done to allow the treatments to diffuse in the CAM.

### Data Gathering Techniques

The duck eggs were cracked carefully in the middle to avoid spillage of the whole content including the embryo and the CAM. The embryo and the CAM were transferred into a sterile petri dish and properly arranged so that the major branching blood vessels and the duck embryo were properly assessed. The anti-angiogenic activity was evaluated using a scoring system of 0-2. The scoring system done is shown in Table 1 below.

**Table 1.** Chorioallantoic Membrane Assay Scoring System.

| Score | Effect | Definition |
| --- | --- | --- |
| 0 | Absent | Normal embryonic development. |
| 0.5 | Weak | There is a little decrease on density of capillary compared to untreated egg. |
| 1 | Moderate | The density of capillary is more than half the normal untreated egg. |
| 2 | Strong | The density of capillary and blood vessels formed is less than half the normal untreated egg. |

After scoring, the following equation was used for the determination of the average mean score for each treatment:

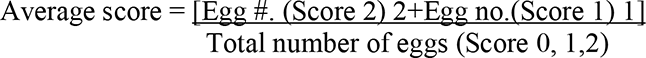

The interpretation of the score is as follows: If the average score is <0.5 this means there is no anti-angiogenic effect. If the average score is 0.5 to 0.75 there is a weak anti-angiogenic effect; If the average score is >0.75 to 1 means there is a good anti-angiogenic effect. The score of >1 means very good anti-angiogenic effect.

### Statistical Treatment

Anti-angiogenic scores of the treatments were analyzed using ANOVA, LSD, and pair-wise comparison among the treatments, respectively. A p-value of less than 0.01 was considered as statistically significant.

## RESULTS AND DISCUSSION

Table 1 showed Treatment 1 (Pure Alugbati Extract) has an average score of 1.33, Treatment 2 (Pure Lumboy Extract) has an average score of 1.67, Retinol palmitate (Positive Control) has an average score 1.78 and Sterilized Normal Saline Solution (Negative Control) has an average score of 0.22. It also depicts the data of each treatment per 3 trials and replicates.

**Table 2.** Anti-angiogenic scores of each treatment.

| Treatments | Trial<sub>1</sub> |  |  | Trial<sub>2</sub> |  |  | Trial<sub>3</sub> |  |  | Ave. | Anti-angiogenic evaluation |
| --- | --- | --- | --- | --- | --- | --- | --- | --- | --- | --- | --- |
|  | R<sub>1</sub> | R<sub>2</sub> | R<sub>3</sub> | R<sub>1</sub> | R<sub>2</sub> | R<sub>3</sub> | R<sub>1</sub> | R<sub>2</sub> | R<sub>3</sub> |  |  |
| T<sub>1</sub>- Pure Alugbati Extract | 2 | 2 | 2 | 2 | 2 | 2 | 1 | 1 | 1 | 1.33 | Very good anti-angiogenic effect |
| T<sub>2</sub>- Pure Lumboy Extract | 1 | 1 | 2 | 1 | 2 | 2 | 1 | 1 | 1 | 1.67 | Very good anti-angiogenic effect |
| T<sub>0+</sub>-Positive Control | 2 | 1 | 2 | 2 | 2 | 2 | 2 | 2 | 1 | 1.78 | Very good anti-angiogenic effect |
| T<sub>0</sub>- Negative Control | 0 | 0 | 1 | 0 | 1 | 0 | 1 | 0 | 1 | 0.22 | No anti-angiogenic effect |

Fig. 1 showed the anti-angiogenic activity of each treatment where retinol palmitate(Positive Control) showed the highest activity followed by Pure Lumboy Extract (Treatment 2), and Pure Alugbati Extract (Treatment 1).This anti-angiogenic capacity of retinol palmitate was already reported in the studies of Fritz, et al., (2011) and Nagpal & Chandraratna (1996). ANOVA revealed that the number of new-formed blood vessel branches among treatments, a 23.806 F-computed value and .000 probability value was attained. The significant difference in the anti-angiogenic activity between treatment 1, treatment 2, and treatment 3 was also tested using LSD. Treatment 1 was compared to treatment 3, treatment 2 to treatment 3 and treatment 1 to treatment 2; these attained the mean differences -0.44444 with 0.039 as significant value; -.011111 with 0.593 as significant value; and -0.33333 with 0.115 as significant value, respectively where all significant values obtained were greater than 0.01 level of significance.

**Figure 1.**
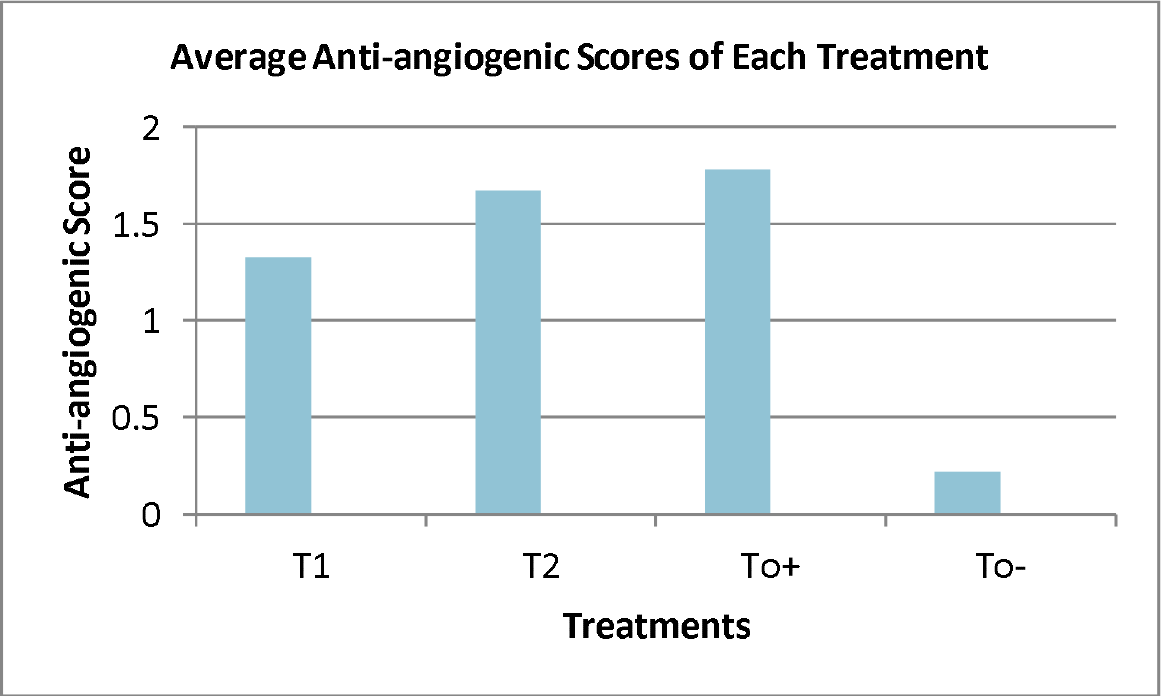
The average anti-angiogenic scores of each treatment.

This computed F-value denotes a significant difference rejecting the null hypothesis of no significant difference among the anti-angiogenic capacity of *B. rubra, S. cumini,* retinol palmitate and sterilized normal saline solution.

The mean difference between *B. rubra* (*Alugbati*) and *S. cumini* extracts compared sterilized normal saline solution has a significant difference. Statistically, the mean of treatments 1 and 2 are higher compared to the negative control which would mean that *S. cumini* and *B. rubra* extracts have anti-angiogenic activity compared to sterilized normal saline solution.

LSD denotes no significant difference between the means of *S. cumini* (Lumboy) and *B. rubra* (Alugbati) extracts as compared to positive control (Retinol palmitate). In addition, the test of significant difference in the anti-angiogenic activity between treatments using Least Significant Difference (LSD) depicts that the mean difference between treatment 1 extract and treatment 2 *S. cumini* (Lumboy) extract has no significant difference. This means that the mean of treatment 2 *S. cumini* (Lumboy) extract is not significantly higher than the mean of treatment 1 *B. rubra* (Alugbati) extract. This implies that *B. rubra* (Alugbati) and *S. cumini* (Lumboy) extracts have equal capacity to inhibit the growth of new blood vessel branches. Furthermore, these findings revealed that both treatments (*B. rubra* (Alugbati) and *S. cumini* (Lumboy) extracts) have compounds that are potential to have anti-angiogenic properties. Both *S. cumini* (Lumboy) and *B. rubra* (Alugbati) extracts have potential anti-angiogenic activity. This could then be further explored for anti-carcinogenic activities.

## RECOMMENDATIONS

It is recommended of using eggs coming from the same hen as treatments for uniform genetic make up. Eggs with different genetic make up might have different angiogenic activity. Also, a toxicity test would be conducted for both *B. rubra* (Alugbati) and *S. cumini* (Lumboy) fruits and other vegetative parts.

